# Chronic, Low-Level Oral Exposure to Marine Toxin, Domoic Acid, Alters Whole Brain Morphometry in Nonhuman Primates

**DOI:** 10.1101/439109

**Authors:** Rebekah Petroff, Todd Richards, Brenda Crouthamel, Noelle McKain, Courtney Stanley, Kimberly S. Grant, Sara Shum, Jing Jing, Nina Isoherranen, Thomas M. Burbacher

**Affiliations:** Department of Environmental and Occupational Health Sciences, University of Washington, Seattle, Washington, USA; Department of Radiology, University of Washington, Seattle, WA, United States; Center on Human Development and Disability, Seattle, Washington, USA; Department of Pharmaceutics, University of Washington, Seattle, Washington, USA; Infant Primate Research Laboratory, Washington National Primate Research Center, Seattle, Washington, USA

**Author notes:** Corresponding Author: Rebekah Petroff Department of Environmental and Occupational Health Sciences School of Public Health, Box 357234 1959 NE Pacific Street University of Washington Seattle, WA 98195.

**Keywords:** Domoic acid, neurotoxicity, diffusion tensor imaging, magnetic resonance spectroscopy, fractional anisotropy, chronic exposure

## Abstract

Domoic acid (DA) is an excitatory neurotoxin produced by marine algae and responsible for Amnesiac Shellfish Poisoning in humans. Current regulatory limits (~0.075-0.1 mg/kg/day) protect against acute toxicity, but recent studies suggest that the chronic consumption of DA below the regulatory limit may produce subtle neurotoxicity in adults, including decrements in memory. As DA-algal blooms are increasing in both severity and frequency, we sought to better understand the effects of chronic DA exposure on reproductive and neurobehavioral endpoints in a preclinical nonhuman primate model. To this end, we initiated a long-term study using adult, female *Macaca fascicularis* monkeys exposed to daily, oral doses of 0.075 or 0.15 mg/kg of DA for a range of 321-381, and 346-554 days, respectively. This time period included a pre-pregnancy, pregnancy, and postpartum period. Throughout these times, trained data collectors observed intentional tremors in some exposed animals during biweekly clinical examinations.

The present study explores the basis of this neurobehavioral finding with in vivo imaging techniques, including diffusion tensor magnetic resonance imaging and spectroscopy. Diffusion tensor analyses revealed that, while DA exposed macaques did not significantly differ from controls, increases in DA-related tremors were negatively correlated with fractional anisotropy, a measure of structural integrity, in the internal capsule, fornix, pons, and corpus callosum. Brain concentrations of lactate, a neurochemical closely linked with astrocytes, were also weakly, but positively associated with tremors. These findings are the first documented results suggesting that chronic oral exposure to DA at concentrations near the current human regulatory limit are related to structural and chemical changes in the adult primate brain.

## 1. INTRODUCTION

Domoic acid (DA) is an excitatory neurotoxin produced by marine algae in the family *Pseudo-nitzschia* and found in ocean waters around the world. DA can accumulate in many types of seafood, including razor clams, scallops, oysters, mussels, anchovies, sardines, and crabs (Andjelkovic et al., 2012; Lefebvre et al., 2002; Trainer et al., 2007; Wekell et al., 1994). When DA-contaminated seafoods are consumed, people may experience symptoms that include gastrointestinal distress, seizures, and the disruption of memory processes, collectively known as the clinical syndrome, Amnesic Shellfish Poisoning (Perl et al., 1990a; Perl et al., 1990b). The largest known DA human poisoning episode occurred in 1987 on Prince Edward Island, Canada, where over 150 people became ill and four died after consuming DA-contaminated mussels. Clinical T2-weighted magnetic resonance (MR) imaging shortly before the death of intoxicated adults displayed stark atrophy of the hippocampus (Cendes et al., 1995). Post-mortem histology in affected patients suggested that DA excitotoxicity was associated with gross necrosis, astrocytosis, and atrophy, primarily in the limbic system and temporal lobe of the brain, including the hippocampus, amygdala, and thalamus (Carpenter, 1990), and similar effects have been documented in a number of model animals and sentinel species after acute DA poisoning (McHuron et al., 2013; Silvagni et al., 2005; Tryphonas et al., 1990; Vieira et al., 2015). Since 1987, there have been no documented cases of human DA poisonings, but toxic algal blooms have been increasing in both severity and frequency (Smith et al., 2018a; Wells et al., 2015). This oceanographic change has been linked to many causal factors, including both seasonal upwelling (Du et al., 2016; Schnetzer et al., 2013; Seubert et al., 2013; Smith et al., 2018b) and shifting ocean temperatures (McCabe et al., 2016; Mckibben et al., 2017; Zhu et al., 2017).

To protect human health, the US Food and Drug Administration has established an action level of 20 ppm in shellfish tissue (US Food and Drug Administration, 2011). This regulatory limit has been officially accepted for commercial and recreational shellfish harvesting in coastal US states (California Office of Health and Environmental Assessment, 1991; Washington Department of Fish and Wildlife, n.d.), as well as in the European Union (O’Mahony, 2018) and Canada (Canadian Food Inspection Agency, 2011). When DA concentrations are at or above 20 ppm in these locations, beaches are closed to shellfish harvesting and commercial fisheries suspend operations (Wekell et al., 2004). The 20 ppm action level was established after the 1987 poisoning, when it was estimated that people showing symptoms of toxicity consumed approximately 200 μg DA. Follow-up studies have calculated that the regulatory limit is equivalent to approximately 0.075-0.10 mg DA/kg bodyweight in a normal, human adult (Mariën, 1996; Toyofuku, 2006). This regulatory limit was, however, only established on acute toxicity data and, in recent years, there has been a growing number of studies documenting the health effects of chronic low-level DA exposure. Data from rodent laboratory research with adult animals suggest that chronic, low-dose exposure can result in short-term, yet recoverable, deficits in cognition (Lefebvre et al., 2017). Human epidemiological findings from a coastal cohort of adult Native Americans in Washington State link the consumption of >15 razor clams/month (a proxy for low-level, chronic DA exposure) to decreased performance on several different memory exams (Grattan et al., 2018, 2016). Cognitive deficits from these epidemiological studies were severe enough to interfere with daily living skills. Collectively, these data suggest that chronic exposure to DA, at environmentally relevant levels of exposure, may have significant consequences on the central nervous system.

One opportunity with which the effects of chronic DA exposure on health and behavior are studied has been sentinel marine species, naturally exposed to DA through the consumption of contaminated seafood (Bossart, 2011). Elevated levels of DA in plasma and urine have been documented in a variety of animals (Lefebvre et al., 2016), but DA toxicity has been most well-defined in California sea lions. Many afflicted animals display symptoms that are similar to those observed in acutely poisoned humans, including changes in cognition, seizures, and, in the case of sea lions, a death rate exceeding 50% (Gulland et al., 2002; Scholin et al., 2000). Sickened animals exhibit signs of gliosis and neuronal necrosis in patterns similar to human DA toxicity cases, with damage primarily in the hippocampus and dentate gyrus (Silvagni et al., 2005). Importantly for the present study, researchers have connected chronic DA toxicosis in sea lions to differences in the structural integrity of the brain, using diffusion tensor imaging (DTI) (Cook et al., 2018). DTI is a model used with diffusion-weighted imaging (DWI), a variation of MR imaging that measures the diffusion rate and anisotropy, or the degree of directionality, of water in tissues. These measures can be used to estimate changes in the density or integrity of axon bundles and myelin, as well as changes in glial cells or extracellular fluids. Cook and colleagues conducted a post-mortem DTI analysis of sea lions diagnosed with DA toxicosis and found decreased anisotropy in the fornix, a white matter tract connecting the hippocampus and thalamus. These data demonstrate a link between oral DA exposure and changes in the microscopic architecture of the mammalian brain, but the translational value of these studies is difficult to ascertain due to differences in neuroanatomy and the lack of quantifiable dose-response data.

The study described in this paper offers an innovative approach to examine the effects of lower level DA exposure by linking behavioral intentional tremors in a nonhuman primate model with in vivo changes in brain structure. Macaques utilized in the present research were selected from a larger, longitudinal reproductive and developmental study (Burbacher et al., in press). In the parent study, adult female macaque monkeys were chronically exposed to 0.0, 0.075 or 0.15 mg/kg/day oral DA prior to, during, and post pregnancy. These exposures were selected to mirror estimates of DA exposure in humans who consumed shellfish with elevated levels of DA below the regulatory threshold (Ferriss et al., 2017; Kumar et al., 2009). Long term exposure in this investigation yielded unanticipated signs of neurotoxicity in the adult females in the form of subtle intentional tremors during a reaching and grasping task (Burbacher et al., in press). Subsequently, the aim of the present imaging study was to explore how the observed intentional tremors in DA-exposed animals were related to changes in brain structure and neurochemistry in vivo. Individuals were selected based on individual tremor and dose status and underwent a single, sedated MR scan with DTI to measure whole brain, voxel-wise diffusion measures. We additionally conducted MR spectroscopy to measure neurochemical concentrations of n-acetyl aspartate (NAA), choline, creatinine, glutamate/glutamine (Glx), and lactate, and captured T1- and T2-weighted images to survey for gross lesions. Results from this translational study represent the first presentation of data that describe in vivo structural changes in nonhuman primates after chronic, oral DA exposure at levels close to real-world human exposures.

## 2. MATERIAL AND METHODS

### 2.1 Study Animals

Macaques for the present study were selected from a larger study aimed at investigating the reproductive and developmental effects of chronic, low-level oral exposure to DA (Burbacher et al., in press). Thirty-two healthy, adult female *Macaca fascicularis* were enrolled in the larger reproductive and developmental study. All animals were housed in the Infant Primate Research Laboratory at the Washington National Primate Research Center, paired with a grooming contact social partner, and allowed unrestricted access to water. Monkeys were fed with Purina High Protein Monkey Diet (St. Louis, MO) biscuits twice a day and provided extensive enrichment (fresh produce, toys, movies/music, and frozen foraging treats). All animal procedure guidelines followed the Animal Welfare Act and the Guide for Care and Use of Laboratory Animals of the National Research Council and protocols were approved by the University of Washington Institutional Animal Care and Use Committee.

Animals were pseudo-randomly assigned to one of three treatment groups: control (n=10), 0.075 (n=11), or 0.15 (n=11) mg/kg/day of DA (BioVectra, Charlottetown, PE, Canada). Blinded testers used positive reinforcement techniques to train macaques to drink from a syringe, complete a battery of clinical assessments to monitor toxicity, and undergo unsedated saphenous blood draws (Burbacher et al., 2004). After training was complete, all experimental procedures were conducted for a 2-month pre-exposure period. During this period, animals were dosed daily with a 5% sucrose solution. Daily, oral exposure to DA was initiated after this 2-month run-in period, and blinded testers orally administered 1 ml of either 0 (n=10), 0.075 (n=11), or 0.15 (n=11) mg/kg of DA in 5% sugar water for at least 2 months. All dosing solutions were quality controlled by measuring DA concentrations via validated LC-MS/MS methods (Shum et al., 2018). After at least two months of exposure, enrolled females were bred with treatment naïve males, and dosing continued throughout breeding and pregnancy. Dosing then was continued postpartum, through the MR imaging.

Plasma DA concentrations were monitored with unsedated blood draws from the saphenous vein, taken 5 hours after dosing. Blood was drawn into sodium heparin tubes and centrifuged at 3,000 *g*. Plasma was separated, stored at −20° C, and analyzed using the methods detailed in Shum et al., 2018. Before pregnancy, average plasma DA concentrations were 1.31 ng/ml for the 0.075 mg/kg/day DA group and 3.42 ng/ml for the 0.15 mg/kg/day DA exposure group. No DA was detected in the vehicle dosed control animals. Twenty-eight females conceived, 9 in the control group, 9 in the 0.075 mg/kg/day DA exposure group, and 10 in the 0.15 mg/kg/day DA exposure group. Mean blood DA concentrations during pregnancy were 0.93 ng/ml and 2.93 ng/ml for the 0.075 and 0.15 mg/kg/day DA exposure groups, respectively.

Throughout the study, general health was monitored daily by clinical staff and weights were recorded weekly. Trained and reliable examiners conducted clinical assessments on all dams at least three times per week. Clinical exams were designed to detect behavioral changes in study animals and included visual orientation and tracking, as well as fine motor and tremor assessments. To assess tremors, blinded testers offered individuals a small treat approximately 6-8 inches from the front of the individual homecage, requiring full extension of the individual’s arm. Testers administered three trials/session, at least three days/week. An animal was considered positive for tremors on any test session if a tester noted the presence of tremors during the reach on at least 2 of the 3 trials. All testers maintained a minimum reliability score of 80% with the primary tester, repeated every 6-8 months. One female in the 0.075 mg/kg/day DA exposure group was dropped from the study after a single breeding due to amenorrhea. In addition, a female in the control group was dropped from the tremor assessment analysis after an MRI revealed a lesion in the temporal lobe (see below for additional details).

Previous reported results from the tremor assessment (Burbacher et al., in press) revealed a significant increase in tremors in the 0.15 mg/kg/day DA exposure group, when tremor increase scores (tremor rates observed over the entire DA exposure period up to delivery minus the tremor rates observed during the pre-exposure period) were compared across DA exposure groups (see Fig. 1). The average tremor increase scores for the 3 DA exposure groups were 5.6% ± 1.4% for the controls, 17.7% ± 4.1% for the 0.075 mg/kg/day DA exposure group, and 30.5% ± 8.3% for the 0.15 mg/kg/day DA exposure group

**Fig. 1:**
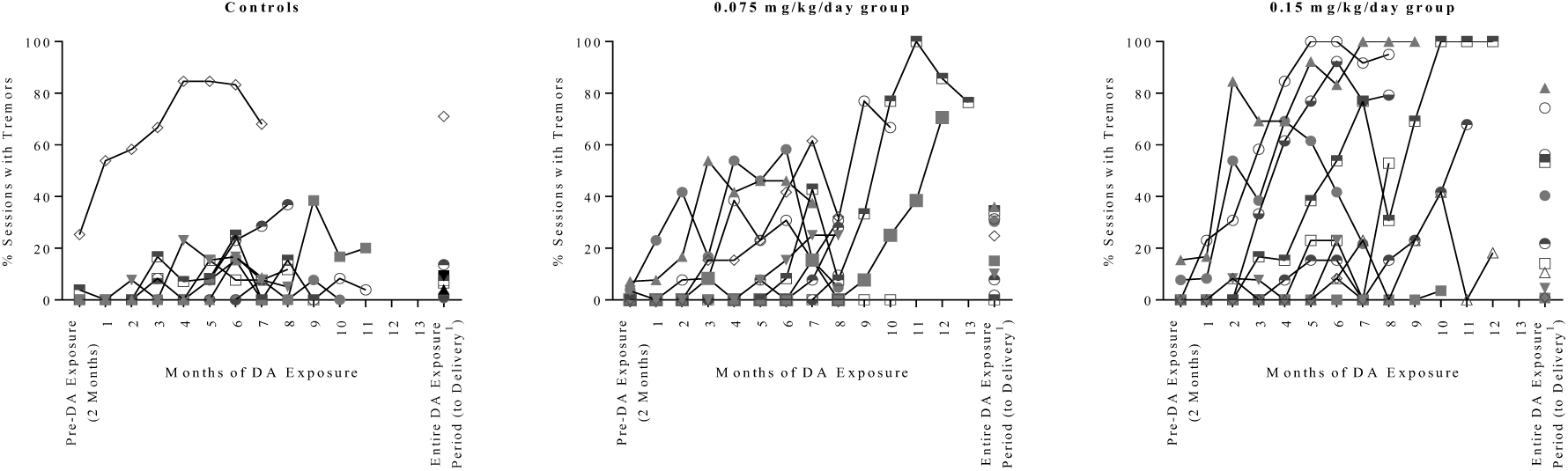
% of Sessions Arm/Hand Tremors Observed on Reaching Task During 2-Month Pre-Exposure Period, Monthly During DA Exposure Period and Over Entire DA Exposure Period to Delivery^1^ (From Burbacher et al., in press)

Enrolled individuals were selected from the larger reproductive and developmental study for MR imaging to compare a subgroup of control females not exhibiting tremors (n=6) to DA exposed females exhibiting tremors (n=6) (Table 1). Selected females included four females from the 0.15 mg/kg/day DA exposure group, two from the 0.075 mg/kg/day DA exposure group and six from the control group. The average age of females selected from the DA exposed and control groups was 8 years and the average weight 4.1 kg. The average duration of DA exposure for the DA exposed females was 419 days.

**Table 1:** Characteristics of Individuals Selected for MRI Study

| ID | Dose (mg/kg/day) | Days Exposed to DA at imaging | Age (years) | Weight (kg) | % Tremors Pre-Exposure<sup>^</sup> | % Tremors Exposure to MRI |
| --- | --- | --- | --- | --- | --- | --- |
| 1 | 0.150 | 363 | 11.58 | 4.80 | 8 | 32 |
| 2 | 0.150 | 546 | 8.08 | 4.30 | 0 | 25 |
| 3 | 0.150 | 554 | 7.94 | 4.10 | 0 | 65 |
| 4 | 0.150 | 346 | 8.27 | 3.05 | 15 | 79 |
| 5 | 0.075 | 381 | 7.96 | 3.95 | 0 | 26 |
| 6 | 0.075 | 321 | 7.52 | 4.40 | 5 | 39 |
| Mean |  | 419 | 8.6 | 4.1 | 5 | 44 |
| 7 | 0.000 | 0 | 9.24 | 5.12 | 0 | 2 |
| 8 | 0.000 | 0 | 7.91 | 3.59 | 0 | 3 |
| 9 | 0.000 | 0 | 7.93 | 3.99 | 0 | 1 |
| 10 | 0.000 | 0 | 8.43 | 5.23 | 3 | 9 |
| 11 | 0.000 | 0 | 8.14 | 4.36 | 0 | 4 |
| 12 | 0.000 | 0 | 7.44 | 3.34 | 0 | 8 |
| Mean |  | 0 | 8.2 | 4.3 | 1 | 5 |
| 13* | 0.000 | 0 | 8.46 | 3.06 | 20 | 64 |

<sup>^</sup> % sessions tremors observed on total sessions tested during a 2-month period immediately preceding the start of exposure.

An additional control female that exhibited a high rate of tremors throughout the study was examined separately to investigate other potential structural brain changes in a non-DA exposed female exhibiting tremors (see Table 1).

### 2.2 MR Image Acquisition and Parameters

Each female underwent a single, sedated MR scan. Less than 30 days before the scan, females were required to meet health standards on a physical exam conducted by clinical veterinary staff. MR image data were acquired on a Philips 3T Achieva (version 5.17) and a custom made 8-channel rf head coil that was developed by Dr. Cecil Hayes and optimized for the small primate head. Females were pre-anesthetized with ketamine (5-10 mg/kg i.m.) and atropine (0.04 mg/kg i.m.) and maintained on inhaled sevoflurane (0.8 - 2.5%) and 100% oxygen. Females were placed in the scanner in prone position, and the coil was arranged over the head. Oxygen saturation levels and single-channel ECG were monitored with an MRI-compatible device (InvivoPrecess^TM^) and temperature was maintained with warm packs.

Diffusion weighted images were acquired with the following parameters: spin-echo echo-planar pulse sequence with diffusion gradients, repetition time 5500 ms, echo time 77.98 ms, reconstructed matrix 128×128, number of slices 44, resolution/voxel size 0.78×0.78×1.5mm, 64 different diffusion weighted directions and one non-diffusion volume at Blip right, b value 1500, 5 different diffusion weighted directions and one non-diffusion volume at Blip left, which where compatible with FSL's topup and eddy software.

Additionally, both a T1-weighted and a T2-weighted image were captured to allow for detection of lesions. The 3-D, high-resolution, T1-weighted MPRAGE images were acquired with a multishot turbo field echo (TFE) pulse sequence and an inversion prepulse (1,151 msdelay); repetition time (TR)/echo time (TE) = 14 s/7.1 ms; 130 axial slices; acquisition matrix 208 x 141 x 130; acquisition voxel size 0.48 x0.53 X1.0 mm; reconstructed voxel size 0.39 x 0.39 x 0.5 mm; slice over sample factor = 2; sense factor = 2 in the foot-head direction; turbo factor = 139; number of signaling averages = 1;TFE shots = 65, and acquisition time = 3 min 14s. A 2-D, high-resolution, T2-weighted images were acquired with a multishot turbo spin-echo (TSE) pulse sequence; repetition time (TR)/echo time (TE) = 7374 ms/80 ms; 24 axial slices; acquisition matrix 208 x 179 x 24; reconstructed voxel size 0.446 x 0.446 x 2 mm; turbo factor = 15; sense factor of 2 in the right left direction, number of signaling averages = 2; and acquisition time = 2 min 42 s.

### 2.3 T1-Weighted and T2-Weighted Image Analysis

Trained testers inspected each slice of T1- and T2-weighted images for abnormalities and lesions in FSLeyes (McCarthy, 2018). Any hypointensities on T1-weighted images and hyperintensities on T2-weighted identified on any single slice were verified as lesions by a second, independent MRI-expert.

### 2.4 Diffusion Weighted Image Processing and Analysis

Whole brain, voxel-wise DTI measures were obtained in FSL (Jenkinson et al., 2012), using a method that is similar to tract-based spatial statistics, but allows for better alignment (Schwarz et al., 2014). Diffusion images were processed using FSL's topup software and FSL's eddy software to minimize distortion from eddy currents and head motion (Andersson et al., 2003; Smith et al., 2004), The FSL program, dtifit (https://fsl.fmrib.ox.ac.uk/fsl/fslwiki/FDT), was used to reconstruct the diffusion tensor for each voxel, and the matrix was diagonalized to obtain tensor eigenvalues, L1, L2, L3. Outcomes of interest included voxel-wise fractional anisotropy (FA), a measure of the directionality of water diffusion and white matter integrity, mean diffusivity (MD, MD=(L1 + L2 + L3)/3), axial diffusivity (AD, AD=L1), and radial diffusivity (RD, RD=(L2+L3)/2). Software buildtemplate, part of Advanced Normalization Tools (ANTs) (Avants et al., 2011), was used to coregister individual FA maps to a target brain, chosen at random from among all scanned individuals. The locations of TFCE significant voxels (from FSL randomise software) were identified using the macaque NeuroMaps atlas (Dubach and Bowden, 2009; Rohlfing et al., 2012).

### 2.5 MR Spectroscopy

MR spectroscopy data were acquired using the same scanner and rf coil described above with the following parameters: PRESS pulse sequence, repetition time 2000 ms, echo time 32 ms, number of FID points 2048, number of averages 48, voxel size 15×15×15 mm and voxel place centered over right thalamus (and including other brain regions) as shown in Fig. 2. MR spectroscopy spar/sdat files were processed using software LCmodel written by Provencher (Provencher, 1993), using both water-suppressed MRS and non-water suppressed MRS files as inputs (Fig. 3). Absolute concentrations of n-acetyl aspartate (NAA), choline, creatinine, glutamate/glutamine (Glx), and lactate were obtained by scaling the in vivo spectrum to the unsuppressed water peak. Concentrations were corrected for cerebral spinal fluid (CSF) volume before statistical analysis.

**Fig. 2.**
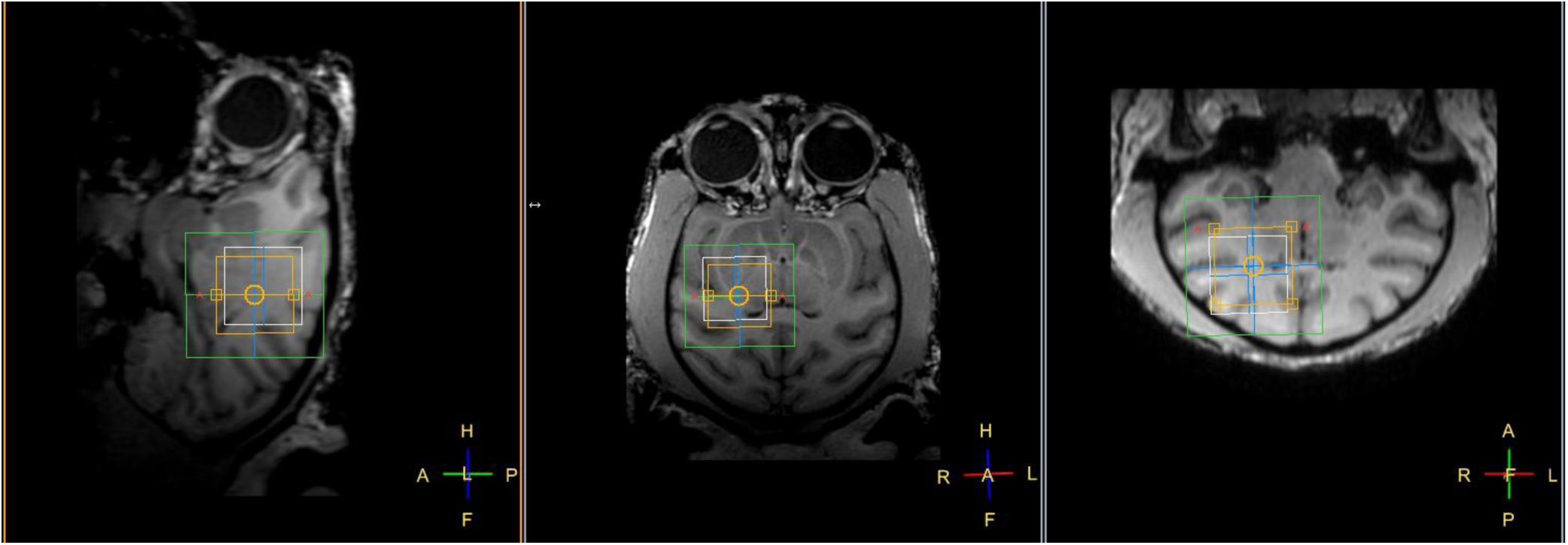
Placement of voxel for MR spectroscopy measurement.

**Fig. 3:**
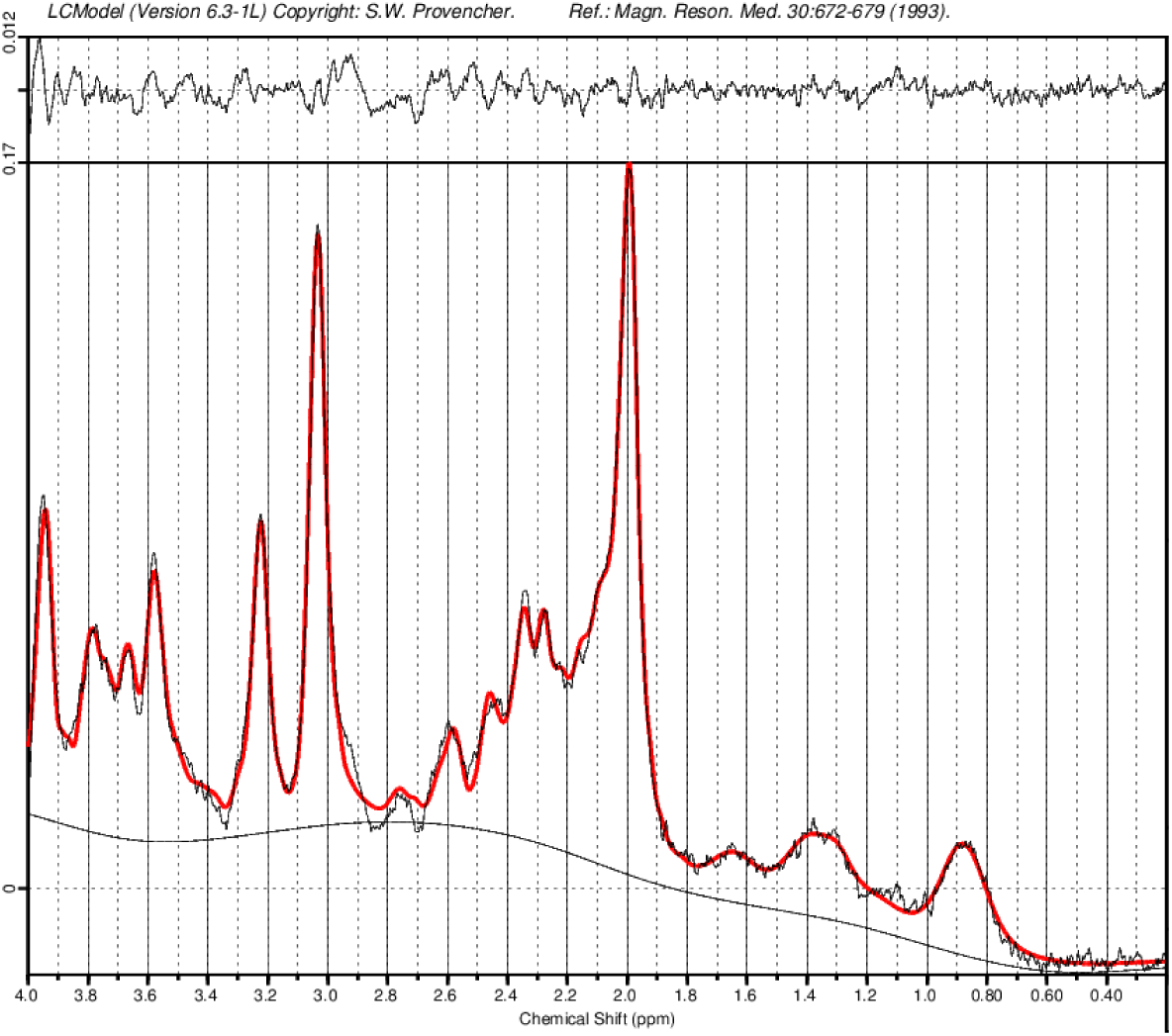
LCmodel fitting of the spectrum from the voxel placement shown in Fig. 2.

**Fig. 4:**
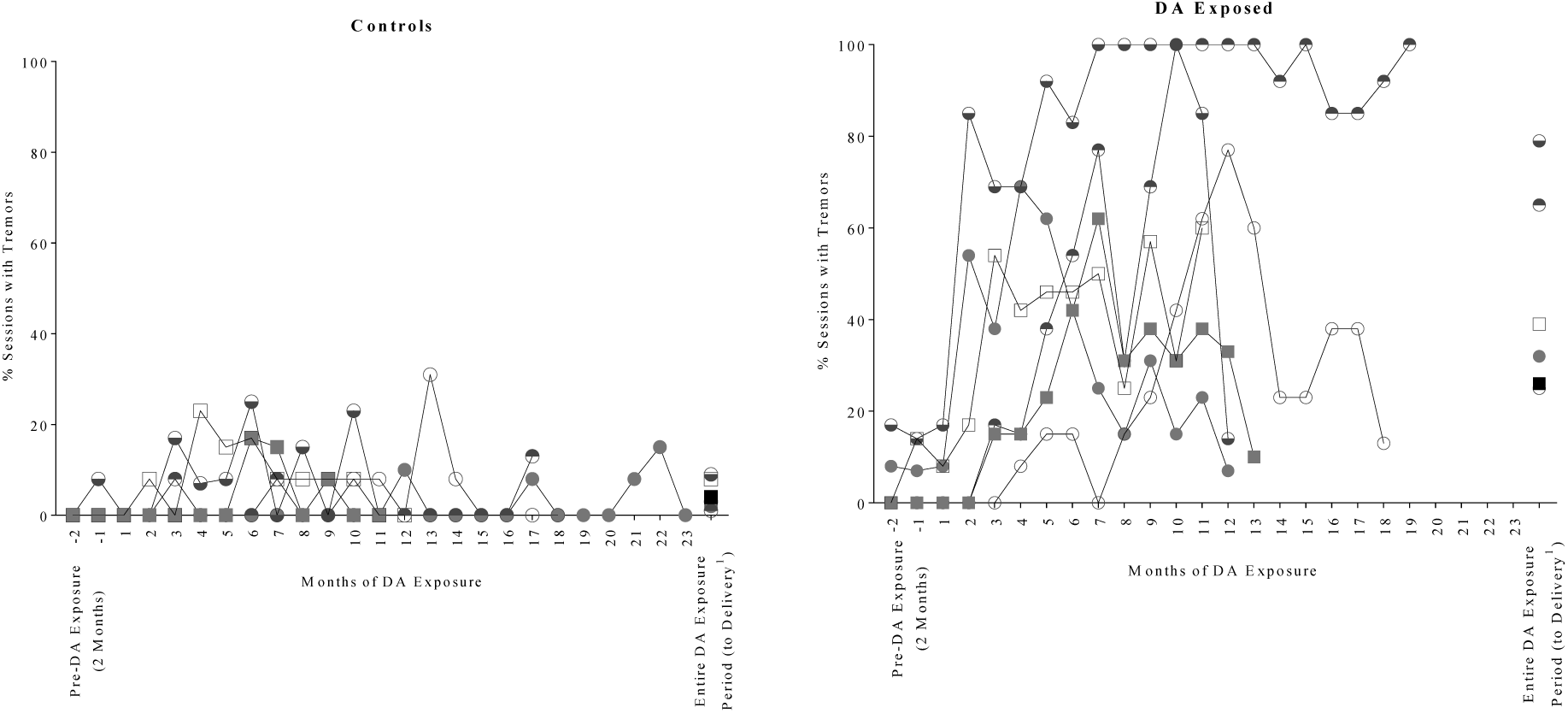
% of sessions arm/hand tremors observed on reaching task monthly during pre-DA exposure period, DA exposure period and over entire DA exposure period to MR study for subset of females selected for MR study.

### 2.6 Statistical Methods

#### 2.6.1 Behavioral Tremors

Tremor scores for the subjects in the present study were calculated during baseline and during the exposure period by dividing the total number of sessions recorded as positive for tremors by the total number of sessions tested. The baseline or pre-exposure period included all clinical sessions from 2 months prior to the first day of exposure through the day before exposure. The exposed tremor score used in all analyses included all clinical observation sessions from day 1 of exposure to the day of imaging. To assess the normality of the exposed tremor scores distribution, a Shapiro-Wilk test was used.

#### 2.6.2 DTI

FSL software randomise, a method that uses 500 random permutations and threshold-free cluster enhancement (TFCE) that corrects for multiple voxel comparisons, was first used to assess group-wise differences between control and exposed groups and then to compute brain-wide correlations of DTI measures and individual tremor score at the time of the scan (Table 1) (Smith and Nichols, 2009; Winkler et al., 2014). Tremor scores were centered/demeaned around the mean tremor score of all animals by subtracting the mean from the individual score, in accordance with the use of this software program. Any significant correlations in either the TFCE randomise software analysis was visually identified in the brain as a cluster in FSLeyes. Diffusion measures from individual voxels within significant clusters were then correlated to the demeaned tremor scores, using the nonparametric Spearman rank method. Because significant clusters were analyzed on a whole-brain level, p-values from the Spearman correlations in individual voxels are not included in the present manuscript.

#### 2.6.3 MRS

Group-wise differences in concentrations of NAA, choline, creatinine, Glx, and lactate were first compared between exposure groups using a Welch’s t-test in R (R Core Team, 2018). A follow-up analysis used Spearman’s correlation in R to assess the correlation between each neurochemical and individual tremor scores (R Core Team, 2018).

### 3. RESULTS

### 3.1 Behavioral Tremors

Tremors were observed rarely during testing sessions prior to DA exposure (see Table 1, average n sessions = 35). The % of sessions that tremors were observed during DA exposure for females selected from the DA exposure groups ranged from 29% to 79%, with an average of 44% ± 9% sessions (n sessions ranged from 117 to 236). The % of sessions that tremors were observed during the DA exposure period for females selected from the control group ranged from 1% to 9%, with an average 4.5% ± 1.3% (n sessions ranged from 134 to 293).

The % of sessions tremors were observed during the DA exposure period up to MR scans (tremor scores) were not normally distributed (W= 0.845, p=0.031), thus nonparametric Spearman’s correlations were used for the analysis of MR measures.

### 3.2 Lesion Identification

Visual inspection of T1- and T2-weighted images revealed that, while there were no lesions in the low-tremoring, controls or high-tremoring, exposed primates (data not shown), the high-tremoring control female (Table 1) had a significant lesion in the right temporal lobe (Fig. 5). Diffusion Tensor Imaging (DTI) and Magnetic Resonance Spectroscopy (MRS) measures for this individual are denoted by a star in Fig. 6 and 7.

**Fig. 5:**
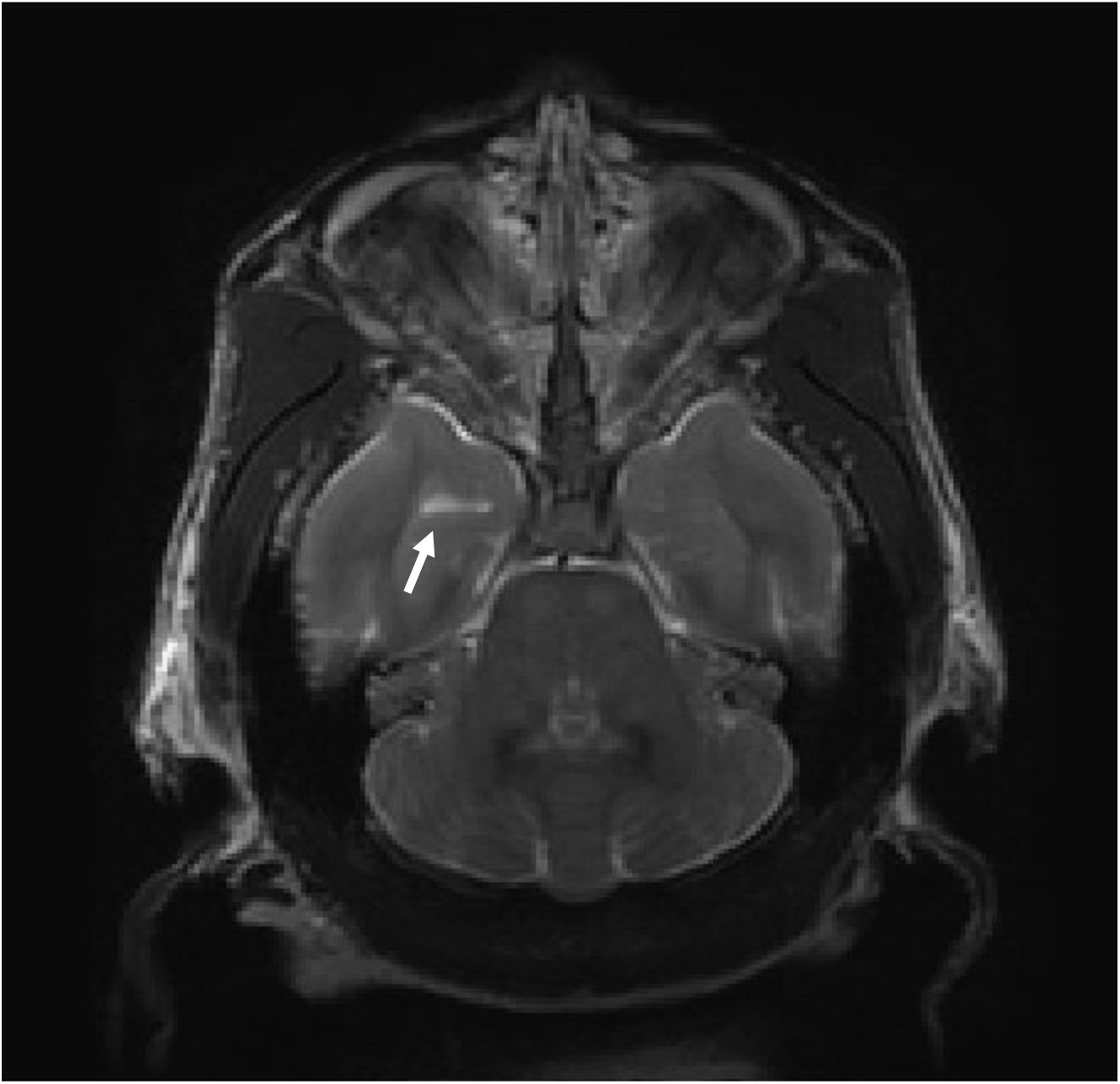
T2-weighted image from the high tremoring, control animal. Lesion indicated with the arrow is located in the right temporal lobe, near the fornix and hippocampus.

**Fig. 6:**
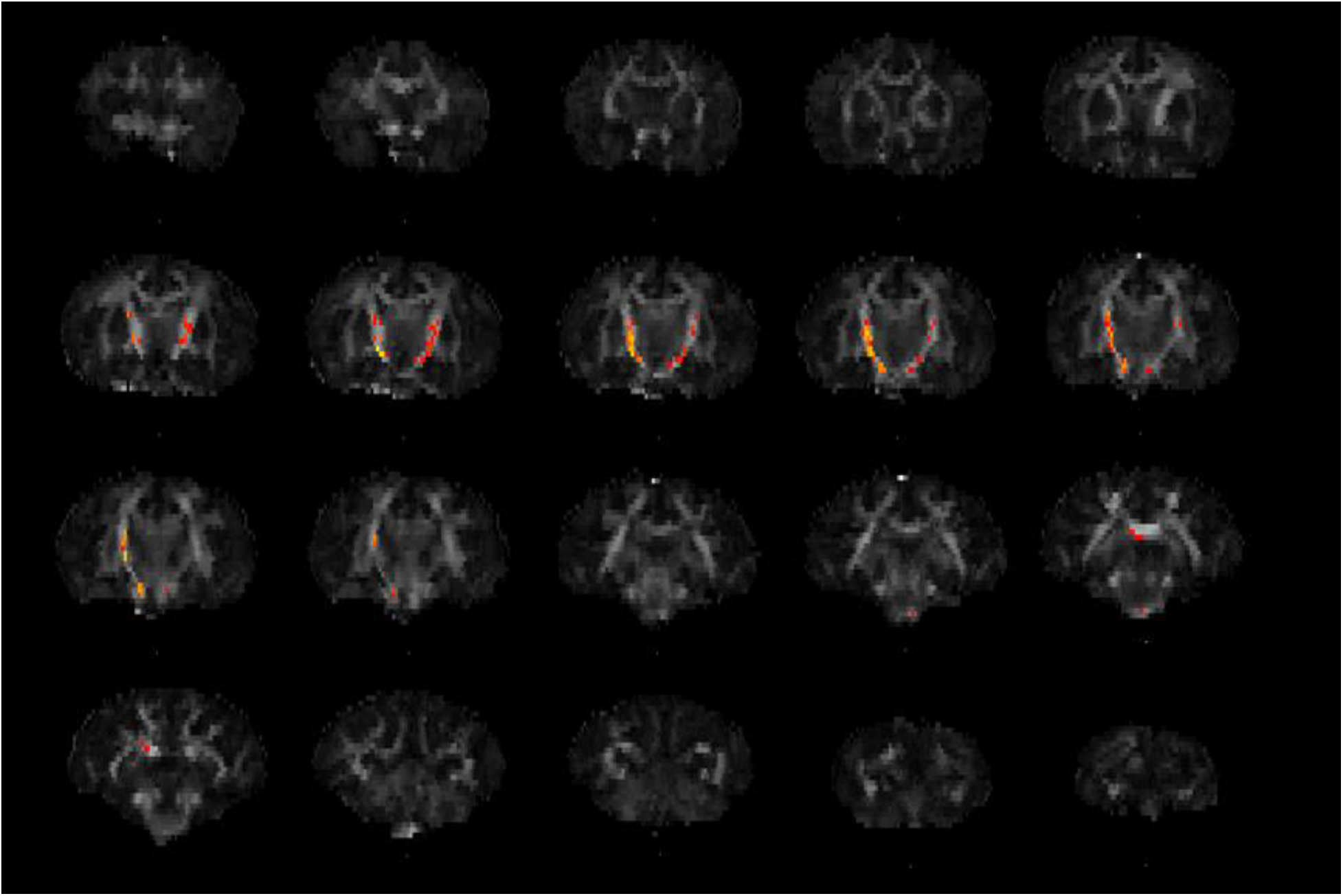
Sequential coronal slices from anterior to posterior, of average FA across all individuals included in the analysis. Significant clusters (p<0.05) are superimposed in red-yellow.

**Fig. 7:**
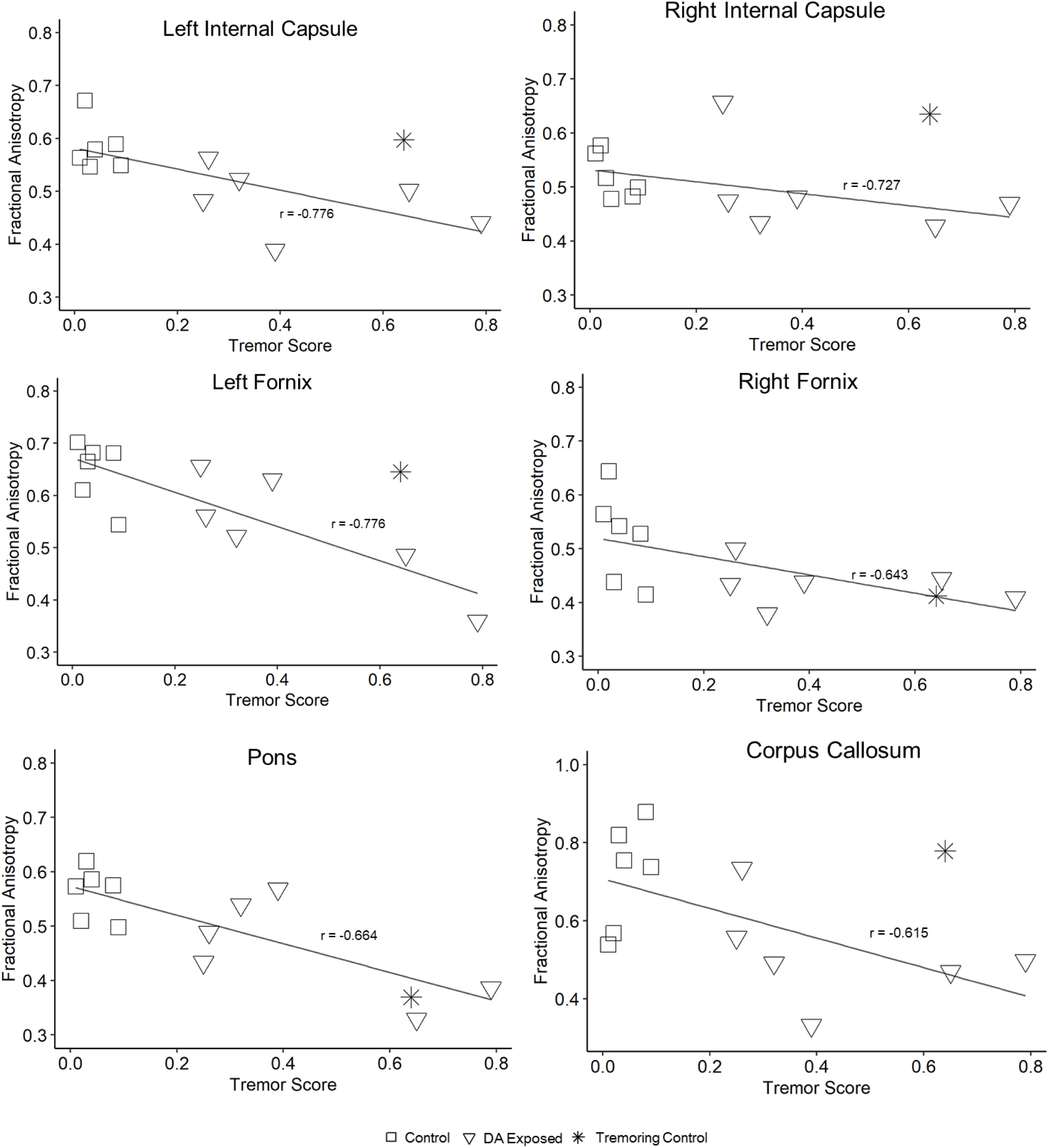
Brain regions with significant Spearman correlations (rho) for tremor scores and FA. Control females are represented by squares, exposed females are shown as triangles. Star denotes the high tremoring control female not included in the correlation analysis.

### 3.3 Diffusion Weighted Images

Using a threshold-free cluster enhancement (TFCE) based analysis, we found that there were no differences in whole-brain DTI measures when using group-wise analysis to compare exposed macaques to controls (fractional anisotropy, p=0.132; axial diffusivity, p=0.392; radial diffusivity, p=0.432; mean diffusivity, p=0.414). Follow-up correlation analysis between whole-brain DTI measures and tremor scores from the 12 scans revealed a significant negative correlation between tremor scores and fractional anisotropy (FA) (p=0.048, Fig. 7). Clusters of FA that were significantly related to tremor scores were observed bilaterally in the anterior internal capsule and fornix. Correlations revealed strong relationships in these regions, as well as with smaller clusters observed in the pons and corpus callosum (Fig. 7). Axial (p=0.178), radial (p=0.218), and mean diffusivity (p=0.232) were not correlated with tremor scores.

### 3.4 MR Spectroscopy

MR spectroscopy concentrations were obtained from each female, centered on the right thalamus. There were no significant differences in neurochemical concentrations between DA exposed and control females (Welch’s t-test; n-acetyl aspartate (NAA), p = 0.924; choline, p= 0.691; creatinine, p=0.086; glutamate/glutamine (Glx), p=0.852; lactate, p=0.908). In addition, CSF-corrected measures for NAA, choline, creatinine, and Glx were not significantly correlated with tremor scores. Lactate concentration, however, was positively correlated with tremor scores, but measurements were highly variable (Fig. 8, p=0.048).

**Fig. 8:**
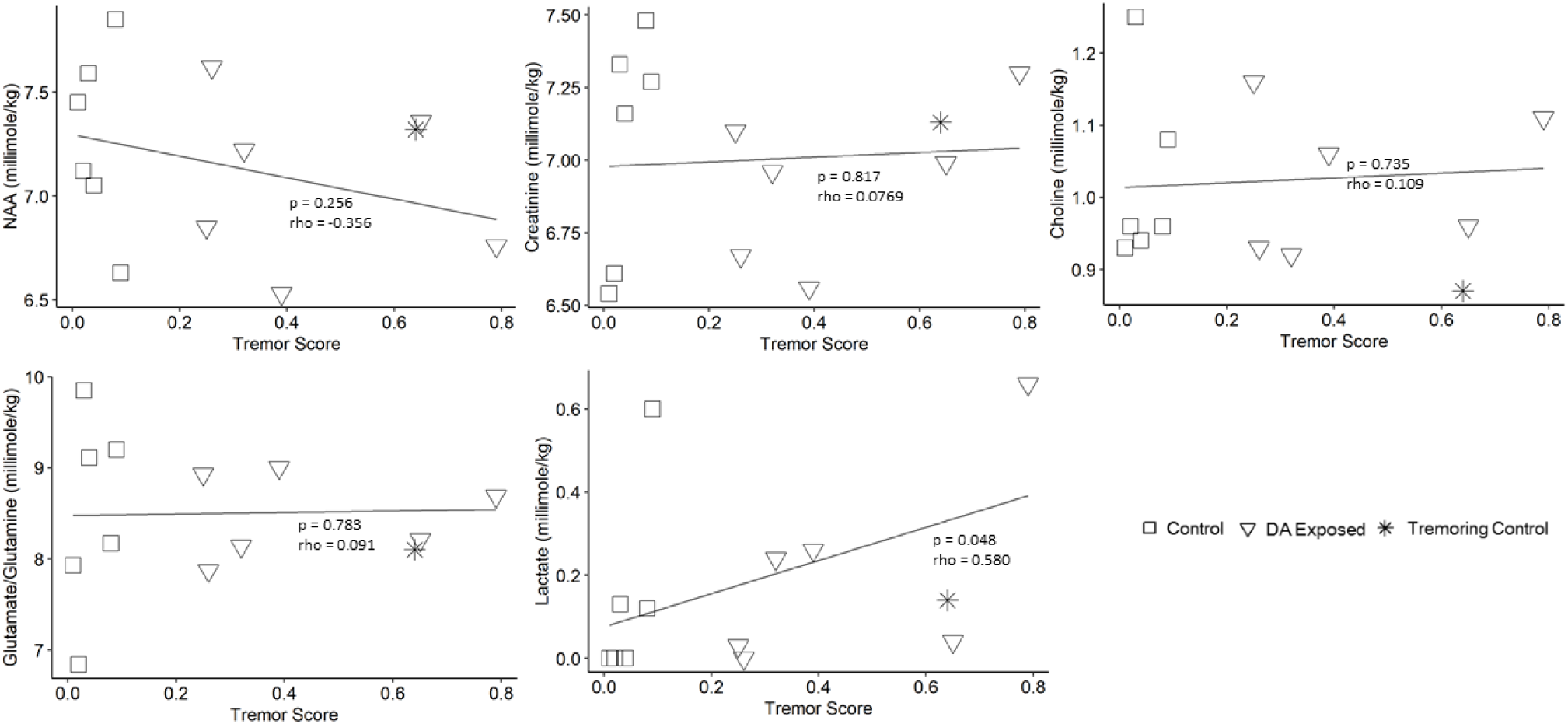
Spearman correlation of CSF-corrected neurochemical concentrations and individual tremor scores. Control females are represented by the squares, exposed females are represented by the triangles. Individual denoted by a star is the high tremoring control female, not included in the analyses.

## 4. DISCUSSION

DA is a known neurotoxin, but few studies have investigated the effects of chronic, low-level exposure to this toxin in any model. This study is the first to use DTI and whole brain analysis in a nonhuman primate model chronically exposed to oral, low-dose DA. We used a TFCE approach with DTI to detect differential clusters of significance, a method that has been shown to have increased sensitivity over other voxel-based analysis methods (Smith and Nichols, 2009) and was untargeted and unbiased. While this approach lowered our ability to detect smaller changes in DTI measures, it allowed us to visualize other structural changes in areas not previously known to be affected by DA. Within the sample of 6 macaque monkeys chronically exposed to low-levels of DA and 6 non-exposed controls, we did not find any changes in DTI measures when comparing DA-exposed macaques to the control group.

However, decreased FA, a measure of tissue integrity, was significantly correlated with increased intentional tremors, but no other diffusion measure was found to be related to tremors. While the fornix, the major white matter tract connecting the hippocampus, the primary target of DA, was affected, there were also other areas of the brain that showed significantly changed FA, including the internal capsule, brainstem, and corpus callosum. Additionally, we found a correlation between tremors and increased lactate in the thalamus. These data collectively show that adult nonhuman primates exposed to chronic, oral, low-levels of DA have neurological damage that can be observed through changes in behavior, neurochemical concentrations, and neurological structural integrity.

Our observed increases in intentional tremors have only been documented in our model, possibly due to the limited number of chronic, oral DA exposure studies. The only other published study to examine chronic oral exposures in a preclinical model used exposure levels of 0.5 and 0.75 mg/kg in macaque monkeys and did not report any significant behavioral or physiological effects after 30 days of repeated exposure (Truelove et al., 1997). It should be further noted that standardized observations, such as those included in the current study, were not utilized in Truelove et al. Other short-term observational and histopathological studies have demonstrated that higher levels of oral exposure (5-10 mg/kg in monkeys and 30-80 mg/kg in rodents) are typically associated with acute symptomology (i.e. scratching, vomiting, shaking/seizures, death) and severe neuronal damage and gliosis primarily in the hippocampus (Iverson et al., 1989; Tryphonas et al., 1990), outcomes not observed in our model. Our results suggest that chronic, low-level oral exposure below levels previously shown to be asymptomatic are related to behavioral tremors.

The present results also suggest that these tremors are connected with decreased FA in several areas of the nonhuman primate brain. FA is a measure of the directionality of water in the brain and ranges in values from 0 (no directionality or equally restricted in all directions) to 1 (fully restricted in one direction). Especially in white matter tracts, FA is typically high and reflects overall axonal integrity (Beaulieu, 2002). It has been suggested that low FA scores are indicative of either direct damage to the myelin/axonal tracts or the replacement of axonal bundles with other cells (i.e. gliosis) (Alba-Ferrara and de Erausquin, 2013; Budde et al., 2011; Garcia-Lazaro et al., 2016; Smith et al., 2006). Significant clusters of decreased FA in tremoring, exposed animals were observed in both the right and left anterior internal capsule and fornix, and smaller clusters were found in the brainstem and the corpus callosum. The internal capsule is a complex bundle of fibers that are essential to motor function (Morecraft et al., 2002), and these fibers include projections that connect the thalamus to the prefrontal cortex, projections from the basal ganglia, and frontopontine fibers that connect the frontal cortex and brain stem (Schmahmann et al., 2004). The pons of the brain stem was also found to have small clusters of decreased FA, possibly in relation to the neurological damage observed in the internal capsule fibers. Clusters of decreased FA were also observed in the fornix, the white matter tract that connects to the hippocampus, the limbic structure responsible for memory and the primary target structure of DA toxicity (Jeffery et al., 2004), and the corpus callosum, the major white matter structure that connects the left and right hemispheres of the brain and is integral to processing stimulation from a multitude of senses (Fabri, 2014). Decreased FA in any of these regions can contribute to a host of neurological phenotypes, but continued research is necessary to understand the underlying connection between the observed behavioral phenotype of intentional tremors to FA deficits across these major brain structures.

In our study, no other diffusion measures, including axial, radial, and mean diffusivity, were changed in relation to tremors. Previous imaging studies have suggested that axial diffusivity reflects acute axonal damage, such as beading (Budde and Frank, 2010) or swelling (Dickson et al., 2007), whereas changes in radial diffusivity are symptomatic of demyelination (Song et al., 2002). Other studies have implicated that when FA is decreased, but mean diffusivity is unchanged, there may be other types of neuronal damage, such as axonal degeneration or an associated glial response, as a cause (Werring et al., 2000). Given our results in FA changes, we propose that there may be axonal degeneration or an increased glial cell response, but not acute axonal damage in primates chronically exposed to low-levels of this increasingly prevalent marine neurotoxin.

Although there are currently no other whole brain DTI analyses in any animal model or human studies of DA exposure, other studies in DA-exposed humans and sea lions have shown that acute DA exposures can produce hippocampal lesions and atrophy as visualized with T2-weighted MR images (Cendes et al., 1995; Montie et al., 2010). Importantly, we did not detect any visible lesions on T1- and T2-weighted images in either the high-tremoring, DA exposed macaques or the low-tremoring control animals, but we did confirm the presence of a lesion on the high-tremoring, control. This finding suggests that tremoring observed in DA exposed animals may be connected with low-level, neurological damage that is not highly visible on T1- or T2-weighted MR images. The only other DTI study conducted in DA-exposed mammals was a post-mortem targeted diffusion analysis in the brains of sea lions that were chronically afflicted with symptoms of DA poisoning (2018). Authors of this MR study used T2-weighted and DTI analyses to assess volumetric and structural changes in the brain. In this study, a limited number brain regions were selected for analysis, and results showed that FA in the fornix, hippocampus, and tracts connecting the hippocampus and thalamus was decreased in DA poisoned animals. These results are similar to those observed in the fornix in our model but were obtained from sea lions demonstrating frank neurotoxicity with visible T2-weighted hippocampal damage, thus suggesting neurological damage that was more severe than the subtle tremors observed in our study. Neurological effects in the fornix, as observed in our study, are also consistent with the published literature, as DA is known to primarily target the hippocampus, resulting in diminished memory. While the present research did not include any examination of cognition, other non-DA, DTI studies in humans have connected decreases in FA to reduced working memory and cognitive performance (Nusbaum et al., 2001; Schulze et al., 2011; Takeuchi et al., 2011).

Our spectroscopy analysis calculated concentrations of several neurochemicals in a voxel placed over the thalamus and adjacent areas of the brain. These data showed that concentrations of NAA, choline, creatinine, lactate and Glx were unchanged in relation to exposure status. NAA, choline, creatinine, and Glx did not correlate with increasing tremors, but lactate was significantly increased with increased tremoring in our cohort. Lactate is an important chemical in the brain, with several multifaceted roles including as: fuel for the brain (Boumezbeur et al., 2010; Smith et al., 2003); signaling for redox cycling and gene expression (Brooks, 2009); and conducting normal astrocyte and myelinating oligodendrocyte functions (Rinholm and Bergersen, 2014). It should be noted, however, that the observed correlation between lactate and tremors was variable (r = 0.58, p=0.048), so further investigations are needed to confirm this.

The neurological damage observed in this study revealed new brain areas that are potential targets of DA, but it should be noted that the present study is exploratory and the first of its kind. Additional research should be conducted in other preclinical models, using both male and female animals, to verify these results and better understand the biochemical and cellular mechanisms underlying the observed changes in FA and lactate. Future research may also be directed at investigating the relationship between FA and DA-related deficits in memory. These results, however, remain compelling for humans who are regularly exposed to DA. Our nonhuman primate model is highly translatable to humans, sharing close similarities in brain structure, connectivity, and function (Passingham, 2009). In addition to our model, we also chose to give exposures orally and near the current regulatory limits (Mariën, 1996; Wekell et al., 2004), to bring strong environmental relevance to the study. These results may be particularly significant to already vulnerable communities that have close cultural connections to various types of seafood, such as some coastal Native American Tribes, where up to 84% of people regularly consume razor clams (Boushey et al., 2016). As DA algal blooms continue to increase in frequency and severity around the globe, it is imperative that we continue to advance our understanding of the health consequences associated with chronic, low-level intake of this marine biotoxin.

## CONFLICT OF INTEREST

None.

## ACKNOWLEDGEMENTS

We would like to thank Mr. Tim Wilbur for this help with the RF coil and MR scanning, staff and volunteers at the Washington National Primate Research Center, and University of Washington Diagnostics Imaging Sciences Center for their skill and technical assistance with this research.

## FUNDING

This research was supported by NIH grants: R01 ES023043, P51 OD010425 and HD083091.

## Footnotes

1 End of breeding for females who did not conceive

## REFERENCES

Alba-Ferrara, L.M., de Erausquin, G.A., 2013. What does anisotropy measure? Insights from increased and decreased anisotropy in selective fiber tracts in schizophrenia. Front. Integr. Neurosci. 7, 9. https://doi.org/10.3389/fnint.2013.00009

Andersson, J.L.R., Skare, S., Ashburner, J., 2003. How to correct susceptibility distortions in spin-echo echo-planar images: Application to diffusion tensor imaging. Neuroimage 20, 870–888. https://doi.org/10.1016/S1053-8119(03)00336-7

Andjelkovic, M., Vandevijvere, S., Van Klaveren, J., Van Oyen, H., Van Loco, J., 2012. Exposure to domoic acid through shellfish consumption in Belgium. Environ. Int. 49, 115–119. https://doi.org/10.1016/j.envint.2012.08.007

Avants, B.B., Tustison, N.J., Song, G., Cook, P.A., Klein, A., Gee, J.C., 2011. A reproducible evaluation of ANTs similarity metric performance in brain image registration. Neuroimage 54, 2033–44. https://doi.org/10.1016/j.neuroimage.2010.09.025

Beaulieu, C., 2002. The basis of anisotropic water diffusion in the nervous system - A technical review. NMR Biomed. https://doi.org/10.1002/nbm.782

Bossart, G.D., 2011. Marine mammals as sentinel species for oceans and human health. Vet. Pathol. 48, 676–690. https://doi.org/10.5670/oceanog.2006.77

Boumezbeur, F., Petersen, K.F., Cline, G.W., Mason, G.F., Behar, K.L., Shulman, G.I., Rothman, D.L., 2010. The contribution of blood lactate to brain energy metabolism in humans measured by dynamic 13C nuclear magnetic resonance spectroscopy. J. Neurosci. 30, 13983–13991. https://doi.org/10.1523/JNEUROSCI.2040-10.2010

Boushey, C.J., Delp, E.J., Ahmad, Z., Wang, Y., Roberts, S.M., Grattan, L., 2016. Dietary assessment of domoic acid exposure: What can be learned from traditional methods and new applications for a technology assisted device. Harmful Algae 57, 51–55. https://doi.org/10.1016/j.hal.2016.03.013

Brooks, G.A., 2009. Cell-cell and intracellular lactate shuttles. J. Physiol. https://doi.org/10.1113/jphysiol.2009.178350

Budde, M.D., Frank, J.A., 2010. Neurite beading is sufficient to decrease the apparent diffusion coefficient after ischemic stroke. Proc. Natl. Acad. Sci. 107, 14472–14477. https://doi.org/10.1073/pnas.1004841107

Budde, M.D., Janes, L., Gold, E., Turtzo, L.C., Frank, J.A., 2011. The contribution of gliosis to diffusion tensor anisotropy and tractography following traumatic brain injury: Validation in the rat using Fourier analysis of stained tissue sections. Brain 134, 2248–2260. https://doi.org/10.1093/brain/awr161

Burbacher, T., Grant, K., Petroff, R., Crouthamel, B., Stanley, C., McKain, N., Shum, S., Jing, J., Isoherranen, N., in press. Effects of Chronic, Oral Domoic Acid Exposure on Maternal Reproduction and Infant Birth Characteristics in a Preclinical Primate Model. Neurotoxicol. Teratol.

Burbacher, T.M., Grant, K.S., Shen, D.D., Sheppard, L., Damian, D., Ellis, S., Liberato, N., 2004. Chronic maternal methanol inhalation in nonhuman primates (Macaca fascicularis): Reproductive performance and birth outcome. Neurotoxicol. Teratol. 26, 639–650. https://doi.org/10.1016/j.ntt.2004.06.001

California Office of Health and Environmental Assessment, 1991. Natural Marine Toxins : PSP and Domoic Acid California’s Mussel Quarantine Natural Marine Toxins : PSP and Domoic Acid Marine Toxin Monitoring Program 1–4.

Canadian Food Inspection Agency, 2011. Canadian Shellfish Sanitation Program: Manual of Operations 1–139.

Carpenter, S., 1990. The Human Neuropathology of Encephalopathic Mussel Toxin Poisoning. Symp. Domoic Acid Toxic. 73–34.

Cendes, F., Andermann, F., Carpenter, S., Zatorre, R.J., Cashman, N.R., 1995. Temporal lobe epilepsy caused by domoic acid intoxication: Evidence for glutamate receptor–mediated excitotoxicity in humans. Ann. Neurol. 37, 123–126. https://doi.org/10.1002/ana.410370125

Cook, P.F., Berns, G.S., Colegrove, K., Johnson, S., Gulland, F., 2018. Postmortem DTI reveals altered hippocampal connectivity in wild sea lions diagnosed with chronic toxicosis from algal exposure. J. Comp. Neurol. 526, 216–228. https://doi.org/10.1002/cne.24317

Dickson, T.C., Chung, R.S., McCormack, G.H., Staal, J.A., Vickers, J.C., 2007. Acute reactive and regenerative changes in mature cortical axons following injury. Neuroreport 18, 283–288. https://doi.org/10.1097/WNR.0b013e3280143cdb

Du, X., Peterson, W., Fisher, J., Hunter, M., Peterson, J., 2016. Initiation and development of a toxic and persistent Pseudo-nitzschia bloom off the Oregon coast in spring/summer2015. PLoS One 11, e0163977. https://doi.org/10.1371/journal.pone.0163977

Dubach, M.F., Bowden, D.M., 2009. BrainInfo online 3D macaque brain atlas: a database in the shape of a brain. Soc. Neurosci. Annu. Meet. Chicago, Abstract No. 199.5.

Fabri, M., 2014. Functional topography of the corpus callosum investigated by DTI and fMRI. World J. Radiol. 6, 895. https://doi.org/10.4329/wjr.v6.i12.895

Ferriss, B.E., Marcinek, D.J., Ayres, D., Borchert, J., Lefebvre, K.A., 2017. Acute and chronic dietary exposure to domoic acid in recreational harvesters: A survey of shellfish consumption behavior. Environ. Int. 101, 70–79. https://doi.org/10.1016/j.envint.2017.01.006

Garcia-Lazaro, H.G., Becerra-Laparra, I., Cortez-Conradis, D., Roldan-Valadez, E., 2016. Global fractional anisotropy and mean diffusivity together with segmented brain volumes assemble a predictive discriminant model for young and elderly healthy brains: A pilot study at 3T. Funct. Neurol. 31, 39–46. https://doi.org/10.11138/FNeur/2016.31.1.039

Grattan, L., Boushey, C.J., Liang, Y., Lefebvre, K.A., Castellon, L.J., Roberts, K.A., Toben, A.C., Morris, J.G.J., Jr., 2018. Repeated Dietary Exposure to Low Levels of Domoic Acid and Problems with Everyday Memory: Research to Public Health Outreach. Toxins (Basel). 10, 103. https://doi.org/10.3390/toxins10030103

Grattan, L., Boushey, C.J., Tracy, K., Trainer, V.L., Roberts, S.M., Schluterman, N., Morris, J.G.J., Jr., Schulterman, N., Morris, J.G.J., 2016. The association between razor clam consumption and memory in the CoASTAL cohort. Harmful Algae 57, 20–25. https://doi.org/10.1016/j.hal.2016.03.011

Gulland, F.M.D., Haulena, M., Fauquier, D., Langlois, G., Lander, M.E., Zabka, T.S., Duerr, R., 2002. Domoic acid toxicity in Californian sea lions (Zalophus californianus): clinical signs. Vet. Rec. 150, 475–480. https://doi.org/doi:10.1136/vr.150.15.475

Iverson, F., Truelove, J., Nera, E., Tryphonas, L., Campbell, J., Lok, E., 1989. Domoic acid poisoning and mussel-associated intoxication: Preliminary investigations into the response of mice and rats to toxic mussel extract. Food Chem. Toxicol. 27, 377–384. https://doi.org/10.1016/0278-6915(89)90143-9

Jeffery, B., Barlow, T., Moizer, K., Paul, S., Boyle, C., 2004. Amnesic shellfish poison. Food Chem. Toxicol. 42, 545–557. https://doi.org/10.1016/j.fct.2003.11.010

Jenkinson, M., Beckmann, C.F., Behrens, T.E.J., Woolrich, M.W., Smith, S.M., 2012. FSL. Neuroimage. https://doi.org/10.1016/j.neuroimage.2011.09.015

Kumar, K.P., Kumar, S.P., Nair, G.A., 2009. Risk Assessment of the amnesiac shellfish poison, domoic acid, on animals and humans. J. Environ. Biol. 30, 319–325. https://doi.org/DOI10.1016/j.automatica.2013.03.005

Lefebvre, K.A., Kendrick, P.S., Ladiges, W., Hiolski, E.M., Ferriss, B.E., Smith, D.R., Marcinek, D.J., 2017. Chronic low-level exposure to the common seafood toxin domoic acid causes cognitive deficits in mice. Harmful Algae 64, 20–29. https://doi.org/10.1016/j.hal.2017.03.003

Lefebvre, K.A., Quakenbush, L., Frame, E., Huntington, K.B., Sheffield, G., Stimmelmayr, R., Bryan, A., Kendrick, P., Ziel, H., Goldstein, T., Snyder, J.A., Gelatt, T., Gulland, F.M.D., Dickerson, B., Gill, V., 2016. Prevalence of algal toxins in Alaskan marine mammals foraging in a changing arctic and subarctic environment. Harmful Algae 55, 13–24. https://doi.org/10.1016/j.hal.2016.01.007

Lefebvre, K.A., Silver, M., Coale, S.L., Tjeerdema, R.S., 2002. Domoic acid in planktivorous fish in relation to toxic Pseudo-nitzschia cell densities. Mar. Biol. 140, 625–631. https://doi.org/10.1007/s00227-001-0713-5

Mariën, K., 1996. Establishing tolerable dungeness crab (Cancer magister) and razor clam (Siliqua patula) domoic acid contaminant levels. Environ. Health Perspect. 104, 1230–6. https://doi.org/10.1289/ehp.961041230

McCabe, R.M., Hickey, B.M., Kudela, R.M., Lefebvre, K.A., Adams, N.G., Bill, B.D., Gulland, F.M.D.D., Thomson, R.E., Cochlan, W.P., Trainer, V.L., 2016. An unprecedented coastwide toxic algal bloom linked to anomalous ocean conditions. Geophys. Res. Lett. 43, 10,366–10,376. https://doi.org/10.1002/2016GL070023

McCarthy, P., 2018. FSLeyes. https://doi.org/10.5281/ZENODO.1887737

McHuron, E.A., Greig, D.J., Colegrove, K.M., Fleetwood, M., Spraker, T., Gulland, F.M., Harvey, J., Lefebvre, K.A., Frame, E.R., 2013. Domoic acid exposure and associated clinical signs and histopathology in Pacific harbor seals (Phoca vitulina richardii). Harmful Algae 23, 28–33. https://doi.org/10.1016/j.hal.2012.12.008

Mckibben, S.M., Peterson, W., Wood, A.M., Trainer, V.L., Hunter, M., White, A.E., 2017. Climatic regulation of the neurotoxin domoic acid. Proc. Natl. Acad. Sci. 114, 239–244. https://doi.org/10.1073/pnas.1606798114

Montie, E.W., Wheeler, E., Pussini, N., Battey, T.W.K., Barakos, J., Dennison, S., Colegrove, K., Gulland, F., 2010. Magnetic resonance imaging quality and volumes of brain structures from live and postmortem imaging of California sea lions with clinical signs of domoic acid toxicosis. Dis. Aquat. Organ. 91, 243–256. https://doi.org/10.3354/dao02259

Morecraft, R.J., Herrick, J.L., Stilwell-Morecraft, K.S., Louie, J.L., Schroeder, C.M., Ottenbacher, J.G., Schoolfield, M.W., 2002. Localization of arm representation in the corona radiata and internal capsule in the non-human primate. Brain 125, 176–198. https://doi.org/10.1093/brain/awf011

Nusbaum, A.O., Tang, C.Y., Buchsbaum, M.S., Tsei Chung Wei, Atlas, S.W., 2001. Regional and global changes in cerebral diffusion with normal aging. Am. J. Neuroradiol. 22, 136–142.

O’Mahony, M., 2018. Eu regulatory risk management of marine biotoxins in the marine bivalve mollusc food-chain. Toxins (Basel). https://doi.org/10.3390/toxins10030118

Passingham, R., 2009. How good is the macaque monkey model of the human brain? Curr. Opin. Neurobiol. https://doi.org/10.1016/j.conb.2009.01.002

Perl, T.M., Bedard, L., Kosatsky, T., Hockin, J.C., Todd, E.C., McNutt, L.A., Remis, R.S., 1990a. Amnesic shellfish poisoning: a new clinical syndrome due to domoic acid. Canada Dis. Wkly. Rep. 16 Suppl 1, 7–8.

Perl, T.M., Bedard, L., Kosatsky, T., Hockin, J.C., Todd, E.C.D., Remis, R.S., 1990b An Outbreak of Toxic Encephalopathy Caused by Eating Mussels Contaminated with Domoic Acid. N. Engl. J. Med. 322, 1775–1780. https://doi.org/10.1056/NEJM199006213222504

Provencher, S.W., 1993. Estimation of metabolite concentrations from localized in vivo proton NMR spectra. Magn. Reson. Med. 30, 672–679. https://doi.org/10.1002/mrm.1910300604

R Core Team, 2018. R: A language and environment for statistical computing. https://doi.org/http://www.R-project.org/

Rinholm, J.E., Bergersen, L.H., 2014. White matter lactate - Does it matter? Neuroscience. https://doi.org/10.1016/j.neuroscience.2013.10.002

Rohlfing, T., Kroenke, C.D., Sullivan, E. V., Dubach, M.F., Bowden, D.M., Grant, K.A., Pfefferbaum, A., 2012. The INIA19 Template and NeuroMaps Atlas for Primate Brain Image Parcellation and Spatial Normalization. Front. Neuroinform. 6, 27. https://doi.org/10.3389/fninf.2012.00027

Schmahmann, J.D., Rosene, D.L., Pandya, D.N., 2004. Motor projections to the basis pontis in rhesus monkey. J. Comp. Neurol. 478, 248–268. https://doi.org/10.1002/cne.20286

Schnetzer, A., Jones, B.H., Schaffner, R.A., Cetinic, I., Fitzpatrick, E., Miller, P.E., Seubert, E.L., Caron, D.A., 2013. Coastal upwelling linked to toxic Pseudo-nitzschia australis blooms in Los Angeles coastal waters, 2005-2007. J. Plankton Res. 35, 1080–1092. https://doi.org/10.1093/plankt/fbt051

Scholin, C.A., Gulland, F.M.D., Doucette, G.J., Benson, S., Busman, M., Chavez, F.P., Cordaro, J., DeLong, R., De Vogelaere, A., Harvey, J., Haulena, M., Lefebvre, K.A., Lipscomb, T., Loscutoff, S., Lowenstine, L.J., Marin, R., Miller, P.E., McLellan, W.A., Moeller, P.D.R., Powell, C.L., Rowles, T.T.K., Silvagni, P., Silver, M.W., Spraker, T., Trainer, V.L., Van Dolah, F.M., 2000. Mortality of sea lions along the central California coast linked to a toxic diatom bloom. Nature 403, 80–84. https://doi.org/10.1038/47481

Schulze, E.T., Geary, E.K., Susmaras, T.M., Paliga, J.T., Maki, P.M., Little, D.M., 2011. Anatomical correlates of age-related working memory declines. J. Aging Res. 2011, 606871. https://doi.org/10.4061/2011/606871

Schwarz, C.G., Reid, R.I., Gunter, J.L., Senjem, M.L., Przybelski, S.A., Zuk, S.M., Whitwell, J.L., Vemuri, P., Josephs, K.A., Kantarci, K., Thompson, P.M., Petersen, R.C., Jack, C.R., 2014. Improved DTI registration allows voxel-based analysis that outperforms Tract-Based Spatial Statistics. Neuroimage 94, 65–78. https://doi.org/10.1016/j.neuroimage.2014.03.026

Seubert, E.L., Gellene, A.G., Howard, M.D.A., Connell, P., Ragan, M., Jones, B.H., Runyan, J., Caron, D.A., 2013. Seasonal and annual dynamics of harmful algae and algal toxins revealed through weekly monitoring at two coastal ocean sites off southern California, USA. Environ. Sci. Pollut. Res. 20, 6878–6895. https://doi.org/10.1007/s11356-012-1420-0

Shum, S., Kirkwood, J.S., Jing, J., Petroff, R., Crouthamel, B., Grant, K.S., Burbacher, T.M., Nelson, W.L., Isoherranen, N., 2018. Validated HPLC-MS/MS Method To Quantify Low Levels of Domoic Acid in Plasma and Urine after Subacute Exposure. ACS Omega 3, 12079–12088. https://doi.org/10.1021/acsomega.8b02115

Silvagni, P.A., Lowenstine, L.J., Spraker, T., Lipscomb, T.P., Gulland, F.M.D.D., 2005. Pathology of domoic acid toxicity in California sea lions (Zalophus californianus). Vet. Pathol. 42, 184–191. https://doi.org/10.1354/vp.42-2-184

Smith, D., Pernet, A., Hallett, W.A., Bingham, E., Marsden, P.K., Amiel, S.A., 2003. Lactate: A preferred fuel for human brain metabolism in vivo. J. Cereb. Blood Flow Metab. 23, 658–664. https://doi.org/10.1097/01.WCB.0000063991.19746.11

Smith, J., Connell, P., Evans, R.H., Gellene, A.G., Howard, M.D.A., Jones, B.H., Kaveggia, S., Palmer, L., Schnetzer, A., Seegers, B.N., Seubert, E.L., Tatters, A.O., Caron, D.A., 2018a. A decade and a half of Pseudo-nitzschia spp. and domoic acid along the coast of southern California. Harmful Algae. https://doi.org/10.1016/j.hal.2018.07.007

Smith, J., Gellene, A.G., Hubbard, K.A., Bowers, H.A., Kudela, R.M., Hayashi, K., Caron, D.A., 2018b. Pseudo-nitzschia species composition varies concurrently with domoic acid concentrations during two different bloom events in the Southern California Bight. J. Plankton Res. 40, 1–17. https://doi.org/10.1093/plankt/fbx069

Smith, S.M., Jenkinson, M., Johansen-Berg, H., Rueckert, D., Nichols, T.E., Mackay, C.E., Watkins, K.E., Ciccarelli, O., Cader, M.Z., Matthews, P.M., Behrens, T.E.J., 2006. Tract based spatial statistics: voxelwise analysis of multi-subjects diffusion data. Neuroimage 31, 1487–1505. https://doi.org/10.1016/j.neuroimage.2006.02.024

Smith, S.M., Jenkinson, M., Woolrich, M.W., Beckmann, C.F., Behrens, T.E.J., Johansen-Berg, H., Bannister, P.R., De Luca, M., Drobnjak, I., Flitney, D.E., Niazy, R.K., Saunders, J., Vickers, J., Zhang, Y., De Stefano, N., Brady, J.M., Matthews, P.M., 2004. Advances in functional and structural MR image analysis and implementation as FSL, in: NeuroImage. Academic Press, pp. S208–S219. https://doi.org/10.1016/j.neuroimage.2004.07.051

Smith, S.M., Nichols, T.E., 2009. Threshold-free cluster enhancement: Addressing problems of smoothing, threshold dependence and localisation in cluster inference. Neuroimage 44, 83–98. https://doi.org/10.1016/j.neuroimage.2008.03.061

Song, S.K., Sun, S.W., Ramsbottom, M.J., Chang, C., Russell, J., Cross, A.H., 2002. Dysmyelination revealed through MRI as increased radial (but unchanged axial) diffusion of water. Neuroimage 17, 1429–1436. https://doi.org/10.1006/nimg.2002.1267

Takeuchi, H., Taki, Y., Sassa, Y., Hashizume, H., Sekiguchi, A., Fukushima, A., Kawashima, R., 2011. Verbal working memory performance correlates with regional white matter structures in the frontoparietal regions. Neuropsychologia 49, 3466–3473. https://doi.org/10.1016/j.neuropsychologia.2011.08.022

Toyofuku, H., 2006. Joint FAO/WHO/IOC activities to provide scientific advice on marine biotoxins (research report). Mar. Pollut. Bull. 52, 1735–1745. https://doi.org/10.1016/j.marpolbul.2006.07.007

Trainer, V., Cochlan, W.P., Erickson, A., Bill, B.D., Cox, F.H., Borchert, J.A., Lefebvre, K.A., 2007. Recent domoic acid closures of shellfish harvest areas in Washington State inland waterways. Harmful Algae 6, 449–459. https://doi.org/10.1016/j.hal.2006.12.001

Truelove, J., Mueller, R., Pulido, O., Martin, L., Fernie, S., Iverson, F., 1997. 30-day oral toxicity study of domoic acid in cynomolgus monkeys: Lack of overt toxicity at doses approaching the acute toxic dose. Nat. Toxins 5, 111–114. https://doi.org/10.1002/nt.5

Tryphonas, L., Truelove, J., Todd, E.C.D., Nera, E., Iverson, F., 1990. Experimental oral toxicity of domoic acid in cynomolgus monkeys (Macaca fascicularis) and rats. Food Chem. Toxicol. 28, 707–715. https://doi.org/10.1016/0278-6915(90)90147-F

US Food and Drug Administration, 2011. Fish and Fishery Products Hazards and Controls Guidance.

Vieira, A.C., Aleman, N., Cifuentes, J.M., Bermudez, R., Pena, M.L., Botana, L.M., Alemañ, N., Cifuentes, J.M., Bermúdez, R., Peña, M.L., Botana, L.M., Aleman, N., Cifuentes, J.M., Bermudez, R., Pena, M.L., Botana, L.M., Alemañ, N., Cifuentes, J.M., Bermúdez, R., Peña, M.L., Botana, L.M., Aleman, N., Cifuentes, J.M., Bermudez, R., Pena, M.L., Botana, L.M., Alemañ, N., Cifuentes, J.M., Bermúdez, R., Peña, M.L., Botana, L.M., 2015. Brain Pathology in Adult Rats Treated With Domoic Acid. Vet. Pathol. 52, 1077–1086. https://doi.org/10.1177/0300985815584074

Washington Department of Fish and Wildlife, n.d. Domoic Acid - A major concern to Washington state’s shellfish lovers [WWW Document]. Fish. Shellfish. URL http://wdfw.wa.gov/fishing/shellfish/razorclams/domoic_acid.html (accessed 1.1.16).

Wekell, J.C., Gauglitz, E.J., Barnett, H.J., Hatfield, C.L., Eklund, M., 1994. The occurrence of domoic acid in razor clams (Siliqua patula), Dungeness crab (Cancer magister), and achovies (Engraulis mordax). J. Shellfish Res. 13, 587–593. https://doi.org/10.2983/035.029.0302

Wekell, J.C., Jurst, J., Lefebvre, K. a, 2004. The origin of the regulatory limits for PSP and ASP toxins in shellfish. J. Shellfish Res. 23, 927–930.

Wells, M.L., Trainer, V.L., Smayda, T.J., Karlson, B.S.O., Trick, C.G., Kudela, R.M., Ishikawa, A., Bernard, S., Wulff, A., Anderson, D.M., Cochlan, W.P., 2015. Harmful algal blooms and climate change: Learning from the past and present to forecast the future. Harmful Algae 49, 68–93. https://doi.org/10.1016/j.hal.2015.07.009

Werring, D.J., Toosy, A.T., Clark, C.A., Parker, G.J.M., Barker, G.J., Miller, D.H., Thompson, A.J., 2000. Diffusion tensor imaging can detect and quantify corticospinal tract degeneration after stroke. J. Neurol. Neurosurg. Psychiatry 69, 269–272. https://doi.org/10.1136/jnnp.69.2.269

Winkler, A.M., Ridgway, G.R., Webster, M.A., Smith, S.M., Nichols, T.E., 2014. Permutation inference for the general linear model. Neuroimage 92, 381–397. https://doi.org/10.1016/j.neuroimage.2014.01.060

Zhu, Z., Qu, P., Fu, F., Tennenbaum, N., Tatters, A.O., Hutchins, D.A., 2017. Understanding the blob bloom: Warming increases toxicity and abundance of the harmful bloom diatom Pseudo-nitzschia in California coastal waters. Harmful Algae 67, 36–43. https://doi.org/10.1016/j.hal.2017.06.004

